# Estimating the benefit of quarantine: eradicating invasive cane toads from islands

**DOI:** 10.1101/344796

**Authors:** Adam S Smart, Reid Tingley, Ben L Phillips

**Author notes:** Corresponding author: Adam Smart, Reid Tingley, Ben Phillips.

## Abstract

1. Islands are increasingly used to protect endangered populations from the negative impacts of invasive species. Quarantine efforts are particularly likely to be undervalued in circumstances where a failure incurs non-economic costs. One approach to ascribe value to such efforts is by modeling the expense of restoring a system to its former state.
2. Using field-based removal experiments on two very different islands off northern Australia separated by > 400 km, we estimate cane toad densities, detection probabilities, and the resulting effort needed to eradicate toads from an island, and use these estimates to examine the financial benefit of cane toad quarantine across offshore islands prioritized for conversation management by the Australian federal government.
3. We calculate density as animals per km of freshwater shoreline, and find striking concordance of density across our two island study sites: a mean density of 353 [286, 446] individual toads per kilometer on one island, and a density of 366 [319, 343] on the second. Detection probability differed between the two islands.
4. Using a removal model and the financial costs incurred during toad removal, we estimate that eradicating cane toads would, on average, cost between $9444 (based on Horan Island; high detectability) and $18093 AUD (Indian Island; low detectability) per km of available freshwater shoreline.
5. Across islands that have been prioritized for conservation benefit within the toads’ predicted range, we provide an estimate of the value of toad quarantine on each island, and estimate the net value of quarantine efforts to be between $27.25 – $52.20 Million AUD. We explore a proposed mainland cane toad containment strategy – to prevent the spread of cane toads into the Pilbara Bioregion, and estimate its potential value to be between $33.79 – $64.74 M AUD.
6. *Synthesis and applications*. We present a modelling framework that can be used to estimate the value of preventative management, via estimating the length and cost of an eradication program. Our analyses suggest that there is substantial economic value in cane toad quarantine efforts across Australian offshore islands and a proposed mainland toad containment strategy.

## Introduction

It is a truth universally acknowledged that an ounce of prevention is worth a pound of cure. In invasive species management, this can be achieved by preventing human-mediated dispersal of non-indigenous species (Chen *et al*. 2018), by conducting routine surveillance programs aimed at early detection (Holden *et al*. 2015), and via translocation of endangered taxa beyond the current or predicted distributions of invaders (Woinarski *et al*. 2014; Legge *et al*. 2018; Moseby 2018) (National Species Management Plan, 2008). Despite such truisms, conservation managers rarely ascribe value to preventative management. Whilst preventative measures are increasingly being adopted to save imperiled taxa (Burns *et al*. 2012; Commonwealth of Australia 2015), without valuation, we risk falling prey to cognitive biases (e.g., immediacy bias), and so routinely commit substantially more money and effort to tactical, “cure” type approaches, than to strategic “prevention”. Quarantine against invasive species is a case in point; vastly more resources are spent controlling the spread and impact of invaders than are spent on preventing their arrival and establishment (Hoffman & Broadhurst 2016).

Quarantine is particularly likely to be undervalued in circumstances in which a failure incurs non-economic costs (e.g., biodiversity loss) (Leung *et al*. 2002) or when costs or damages persist over long-time scales (Epanchin-Neill *et al*. 2015). One way to place a value on such quarantine efforts is to calculate the cost of restoring the system to its former state (Kimball *et al*. 2014; Rohr *et al*. 2016). In the case of an invasive species with primarily non-economic impacts, we can calculate the ongoing benefit of quarantine as the expense of restoring the system to this former state, i.e., a subsequent eradication program. Such a valuation is a lower bound on the benefit of quarantine for a number of reasons. First, the same quarantine effort typically protects against many potential invasive species. In addition, any impact that an invasive species has before it is eradicated (e.g., local extinction or shifts in population structure of a native species, altered landscape vegetation profiles formation) must be added to the cost of restoration (Hoffmann & Broadhurst 2016, Jardine & Sanchirico 2018). Thus, the cost of eradicating a single invader is a very conservative estimate of the true value of quarantine efforts.

Islands are important resources for conservation quarantine because they offer a natural barrier to the spread of invasive species. Conservation biologists routinely exploit this property of islands, not only to protect species that naturally occur on islands, but also to provide refuge for species under threat on the mainland (Thomas 2011; Tershy *et al*. 2015; Legge *et al*. 2018). In Australia alone, a minimum of 47 conservation translocations to islands have been carried out to date (Department of the Environment, Water, Heritage and the Arts, 2009). In these circumstances – where the conservation value of an island has been artificially bolstered – the subsequent arrival of invasive species can have a larger impact than they otherwise would. Typically, island quarantine is used by conservation managers to protect native species from invasive predators (e.g., foxes, cats, weasels, rats). In Australia, however, islands are also used to mitigate the impact of cane toads (*Rhinella marina*) on native predators (Moro *et al*. 2018; Ringma *et al*. 2018). Cane toads were introduced to northeastern Australia in the 1930s and, in northern Australia, continue to spread westerly at a rate of ~50 km per year (Phillips *et al*. 2010). This invasion has had major impacts on populations of native predators, many of which have no resistance to the toad’s toxin (Nelson *et al*. 2010; Greenlees *et al*. 2010; Llewelyn *et al*. 2014). In response to declines of multiple predator species (e.g., dasyurids, monitors, snakes) the Australian government implemented the Cane Toad Threat Abatement Plan (2011), which aimed to identify, and where possible reduce, the impact of cane toads on native species (Shanmuganathan *et al*. 2010). A lack of viable methods for broad-scale control, however, has since led the Australian government to place an increased emphasis on containment (on the mainland) and on quarantine (on offshore islands) to mitigate the biodiversity impacts of cane toads.

While quarantine is currently the best available strategy, it is not a panacea: cane toads have already established on at least 48 islands across northern Australia (McKinney *et al*. 2018 unpub data), with potential for further natural and anthropogenic introductions. Thus, execution of the strategy outlined in the Cane Toad Threat Abatement Plan requires ongoing quarantine, eradication and containment efforts. Here we estimate the lower bound of the monetary value of these ongoing efforts, by quantifying the cost of eradicating cane toads from two islands in northern Australia. We approach this problem by estimating the density and detection probability of toads on each island, and use these estimates to calculate the amount of time and money it would take to remove enough toads to ensure eradication.

## Materials and methods

### Study Area

This study was carried out on two islands in northern Australia: Horan Island on Lake Argyle, Western Australia (HI) and Indian Island in the Northern Territory (II). Lake Argyle is Western Australia’s largest man-made reservoir covering > 880 km$ and is located within the East Kimberly region. The study site is composed of exposed spinifex-covered hilltops and sparse savanna woodland. Freshwater is available year-round, with the lake contracting from May–November. Toads are thought to have colonized islands on the lake in the wet seasons of 2009/2010 (Somaweera & Shine 2012). Indian Island is an offshore island, 40 km west of Darwin. It supports predominantly savanna woodland and monsoonal vine thicket, with a large ephemeral freshwater swamp located on the northern tip of the island. Depending of the magnitude of the wet season, standing water can be present in this swamp year-round or dry up by late September. Toads are thought to have colonized Indian Island via rafting events around 2008. Access to Indian Island was granted by Kenbi Traditional Owners (Northern Land Council permit 82368).

### Field sampling

Cane toad surveys occurred over six nights, on each island, denoted, *t* = {0,1,…, 5}, during November 2017 (HI) and October 2018 (II). Surveys commenced at sundown each evening and lasted three hours, with ambient temperatures ranging from 24 – 35°C. As Horan Island occurs on a freshwater lake, the entire island was circumambulated each night by two people using headtorches; one individual focused on the higher part of the shoreline, the other on the lower shoreline. Indian Island is an oceanic island, with only a single freshwater swamp present in the dry season. This swamp was navigated each night by two people using head torches. On both islands, every toad encountered was collected and humanely killed on site in accordance with The University of Melbourne animal ethics protocol (1714277.1) and State laws regarding handling of non-native species. Each night, we recorded the number of individuals collected, *c_t_*. Surveys were conducted immediately prior to the breeding season so that only post-metamorphic age classes were encountered.

### Statistical analysis

We do not encounter every individual on a given night, and so incorporate imperfect detection. For each island, we aim to estimate three parameters: *N*_0_, the true number of toads on the island at the commencement of surveys; *p*, the mean per-individual detection probability; and *α*, the length of time (in days) required to eradicate toads from our treatment areas. The number of individuals collected each night, *c_t_*, can be considered a draw from a binomial distribution with:

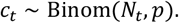

Where *N*_0_, the pre-sampling population size, is a latent variable with a mean and variance equal to λ, such that:

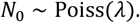

For *t* > 0:

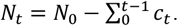

The length of time required to remove a population, *α*, from a treatment area is described via the relationship:

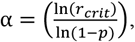

where, *r_crit_*, the critical removal threshold, is equal to 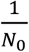 (the inverse of the pre-sampling population size).

Models were fit with Markov chain Monte Carlo (MCMC) in JAGS v.4.6.0, run through R v3.4.1 via the package rjags v4.6.0 (Plummer & Martyn 2013). Three model chains were run for 30,000 iterations, with the first 10,000 iterations discarded as a burn-in, which was sufficient for the MCMC chains to converge. Convergence was checked using the Gelman-Rubin diagnostic (Gelman & Rubin 1992); all chains produced potential scale reduction factors < 1.1, indicating convergence of chains. The remaining samples were thinned by a factor of 2, resulting in 10,000 samples per chain for post-processing. Minimally informative prior distributions for *p* and *»* were specified as uniform between 0 - 1 and 0 - 10,000 respectively.

We denote a successful eradication to have occurred when only a single toad remains (i.e., no further breeding pairs remain). As we assume that removal efforts take place on consecutive nights until completion, we disregard breeding and immigration.

### Cost analysis

We estimate the cost of eradicating toads on our study islands based on consumable, personnel, and travel costs incurred during toad collection (see Appendix S1 in Supporting Information). Relative to most islands across northern Australia, both Horan and Indian Islands are readily accessible, thus our travel costs are modest. We assume that eradication is conducted by a fully-equipped organization; thus we do not include vehicle/boat purchase or hire (i.e., set-up costs), nor do we consider organizational in-kind associated with utilizing existing capital. Removal efforts are carried out on subsequent nights until eradication is reached; therefore, the cost associated with travel to and from our site is incurred only once. Travel costs include a $85/hour consultant rate plus the additional costs of fuel, insurance, and vehicle maintenance (an extra $36/hour). Thus, total travel costs are $111/hour.

### Cost Scenarios

We use our estimates of toad removal on Horan and Indian Islands (with their attendant detection probabilities) to highlight the potential benefit of quarantine efforts on a subset of high priority islands (Table 1). Our chosen islands are drawn from a list of 100 oceanic islands that the Australian Commonwealth has prioritized for conservation, due to their biodiversity value and presence of species listed under the Environment Protection and Biodiversity Conservation Act (Department of the Environment and Energy [DEE], 1999). We refine this list to include only islands that are ≥2 km from the Australian mainland and occur within the potential distribution of cane toads in Australia (Kearney *et al*. 2008). For each island in our dataset, we map the length of permanent freshwater shoreline available, using either satellite maps, government/landholder records, or a combination of both – resulting in a net kilometer length of shoreline for each island in our dataset. All islands were crossed-checked for the presence of cane toads via the ‘Feral Animals on Offshore Islands’ database (DEE, 2016) in addition to the presence of human settlement. In cases where islands had no permanent freshwater but did have human settlement (or known livestock presence), a one-kilometer circumference was assumed around dwellings and visible watering points.

**Table 1:** Islands included in analyses from the top 100 islands prioritized by the Australian Commonwealth for conservation actions (Department of the Environment, Water, Heritage and the Arts (2009)). Estimates for the benefit of quarantine are in ‘000s (AUD). Mean benefit reports the cost of removal, averaging over costs calculated with the detection probabilities of each of our island systems.

| Jurisdiction | Island Name | Toads Present | Distance to mainland (km) | Area (km<sup>2</sup>) | Length of freshwater shoreline (km) | Mean benefit of quarantine (000s) | Lower Est. | Upper Est. |
| --- | --- | --- | --- | --- | --- | --- | --- | --- |
| New South Wales | Lord Howe Island | No | 570 | 11 | 1 | 18 | 10 | 28 |
| Western Australia | Barrow Island | No | 56 | 139 | 21 | 373 | 200 | 580 |
|  | Bernier Island | No | 38 | 171 | 2 | 36 | 19 | 55 |
|  | East Intercourse Island | No | 5.5 | 51 | 2 | 36 | 19 | 55 |
|  | Faure Island | No | 6.1 | 8 | 2 | 36 | 19 | 55 |
| Queensland | Badu Island | Yes | 90 | 53 | 10 | 178 | 95 | 276 |
|  | Bentineck Island | Yes | 25 | 269 | 5 | 89 | 48 | 138 |
|  | Boigu Island | Yes | 7.8 | 6 | 55 | 977 | 524 | 1519 |
|  | Darnley Island | Yes | 70 | 195 | 0 | 18 | 10 | 28 |
|  | Dunk Island | Yes | 4 | 170 | 1 | 18 | 10 | 28 |
|  | Goold Island | Yes | 15 | 101 | 1 | 18 | 10 | 28 |
|  | Hammond Island | Yes | 18 | 104 | 3 | 53 | 29 | 83 |
|  | Horn Island | Yes | 16.7 | 396 | 8 | 142 | 76 | 221 |
|  | Macleay Island | Yes | 3 | 16 | 0.7 | 12 | 7 | 19 |
|  | Magnetic Island | Yes | 6.3 | 6 | 2 | 36 | 19 | 55 |
|  | Moa Island | Yes | 52 | 72 | 21 | 373 | 200 | 580 |
|  | Moreton Island | Yes | 20 | 7 | 54 | 959 | 514 | 1491 |
|  | Mornington Island | Yes | 29 | 1662 | 102 | 1812 | 971 | 2817 |
|  | North Stradbroke Island | Yes | 3.8 | 1001 | 105 | 1865 | 1000 | 2900 |
|  | Prince of Wales Island | Yes | 16 | 148 | 27 | 480 | 257 | 746 |
|  | Sweers Island | No | 30 | 7 | 4 | 71 | 38 | 110 |
| Northern Territory | Bathurst Island | No | 61 | 235 | 137 | 2434 | 1305 | 3783 |
|  | Centre Island | Yes | 7.8 | 64 | 20 | 355 | 190 | 552 |
|  | Croker Island | No | 3 | 11 | 152 | 2700 | 1447 | 4197 |
|  | Groote Eylandt | No | 45 | 42 | 203 | 3606 | 1933 | 5606 |
|  | Marchinbar Island | No | 21 | 5 | 59 | 1048 | 562 | 1629 |
|  | Melville Island | No | 24 | 2 | 1054 | 18724 | 10036 | 29106 |
|  | North Island | Yes | 28 | 13 | 3 | 53 | 29 | 83 |
|  | Peron Island | No | 3.4 | 3 | 3 | 53 | 29 | 83 |
|  | Raragala Island | No | 36 | 52 | 11 | 195 | 105 | 304 |
|  | Vanderlin Island | Yes | 7 | 6 | 68 | 1208 | 647 | 1878 |
|  | West Island | Yes | 4 | 576 | 30 | 533 | 286 | 828 |
|  | Yabooma Island | No | 2.7 | 2 | 3 | 53 | 29 | 83 |

In addition to the islands derived from this report, we explore the value of a potential cane toad containment strategy outlined in a revised version of the Cane Toad Threat Abatement Plan (Tingley *et al*. 2013).

This strategy aims to develop a ‘waterless barrier’ on the Australian mainland by excluding cane toads from artificial water bodies on cattle stations between Broome and Port Hedland in Western Australia. Using a dataset containing the presence of bore holes, cattle watering points, dams and permanent freshwater bodies in the Pilbara bioregion (see Southwell *et al*. 2017) we estimate the economic benefit of the proposed barrier. A one-kilometer circumference was applied to all waterpoints, dams and pools, in addition to a per-kilometer of shoreline rate along permanent watercourses within the region. If implemented successfully, this strategy could keep toads out of the Pilbara (and subsequent regions) – an effective quarantine of 268,00km$ of the Australian mainland (see Florance *et al*. 2011; Tingley *et al*. 2013; Southwell *et al*. 2017 for further information).

## Results

The number of cane toads removed from both Horan and Indian Island, *c_t_*, declined over time (Figure 1). Across the duration of our surveys, we captured and removed a total of 1550 cane toads (1251 on HI, 299 on II). The estimated probability of detecting an individual toad on a given night differed between our two study sites (Horan Island: mean *p* [95% credible interval] = 0.1 [0.07, 0.13]; Indian Island: 0.27 [0.22, 0.33]) (Figure 2). Given the site-specific detection probability, the estimated number of toads present at the initiation of our surveys (*N*_0_) was much higher on Horan Island (2681 [2171, 3393]) than on Indian Island (353 [308, 408]) (Figure 3).

**Figure 1.**
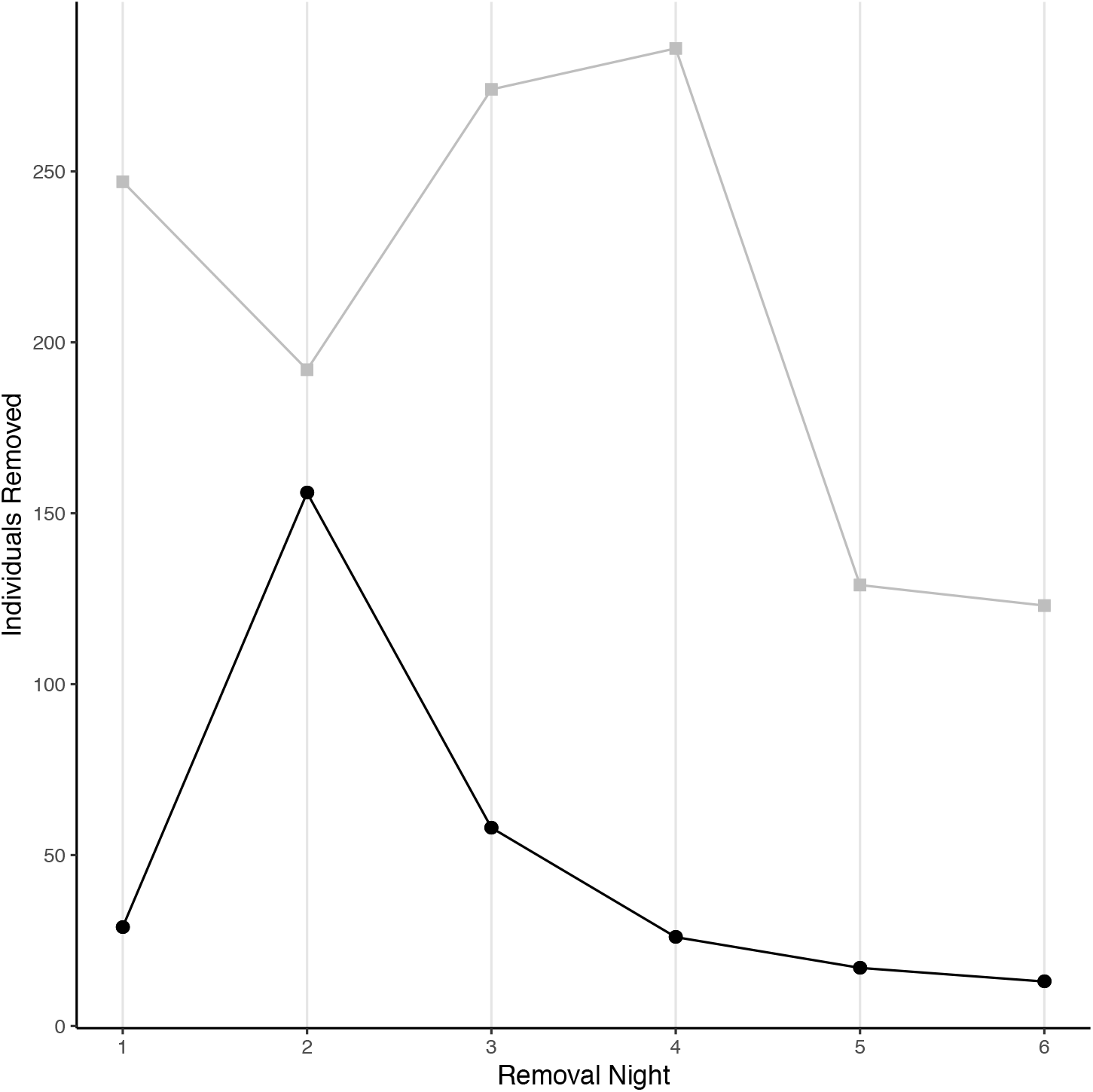
Numbers of individual cane toads captured per night on Horan (gray) and Indian (black) Islands.

**Figure 2.**
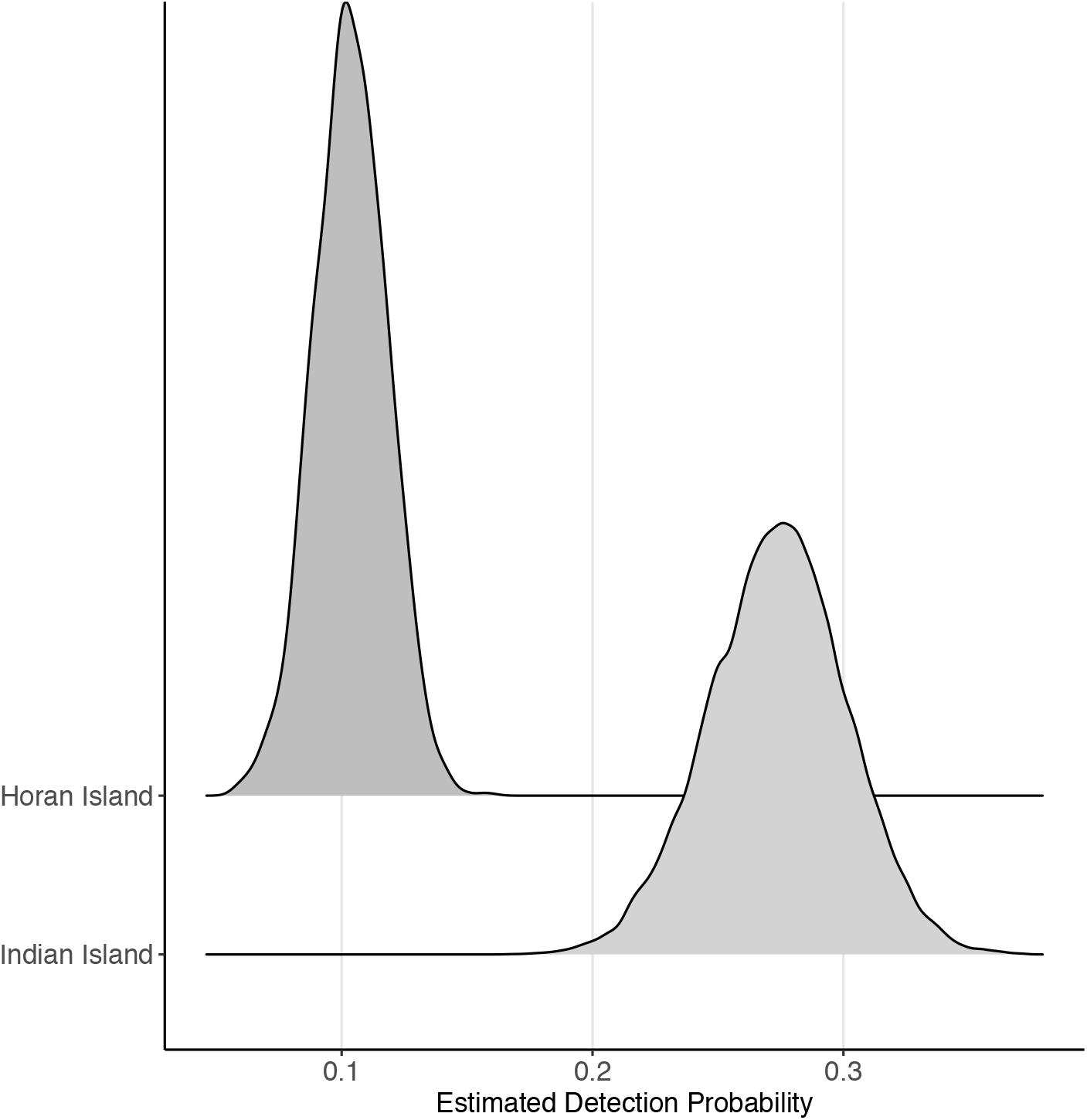
Distributions of the estimated detection probabilities of cane toads on Horan and Indian Islands.

**Figure 3.**
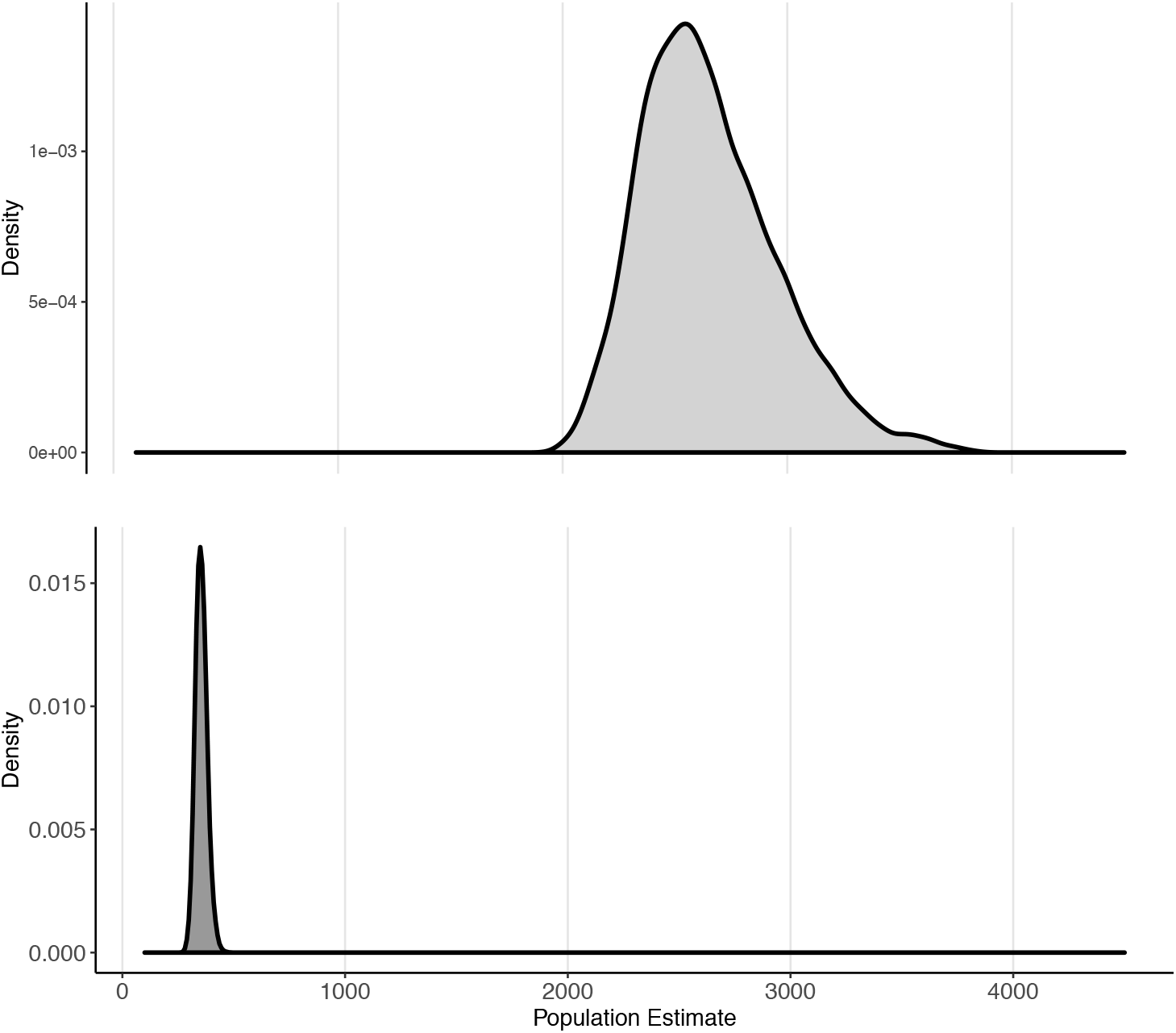
Estimated distributions of cane toad population size estimates (*N*_0_,) before removal efforts.

Horan Island – situated in a freshwater lake – has a circumference of 7.63 km, which translates to a cane toad density of 353 [286, 446] individuals per kilometer of freshwater shoreline. The freshwater source on Indian Island has a circumference of 1.04 km, translating to a density of 366 [319, 343] individuals per kilometer of shoreline (Figure 4). We could also express toad density as animals per km$ of island, in which case we calculate a density of individuals of 56 on II and 2852 on HI.

**Figure 4.**
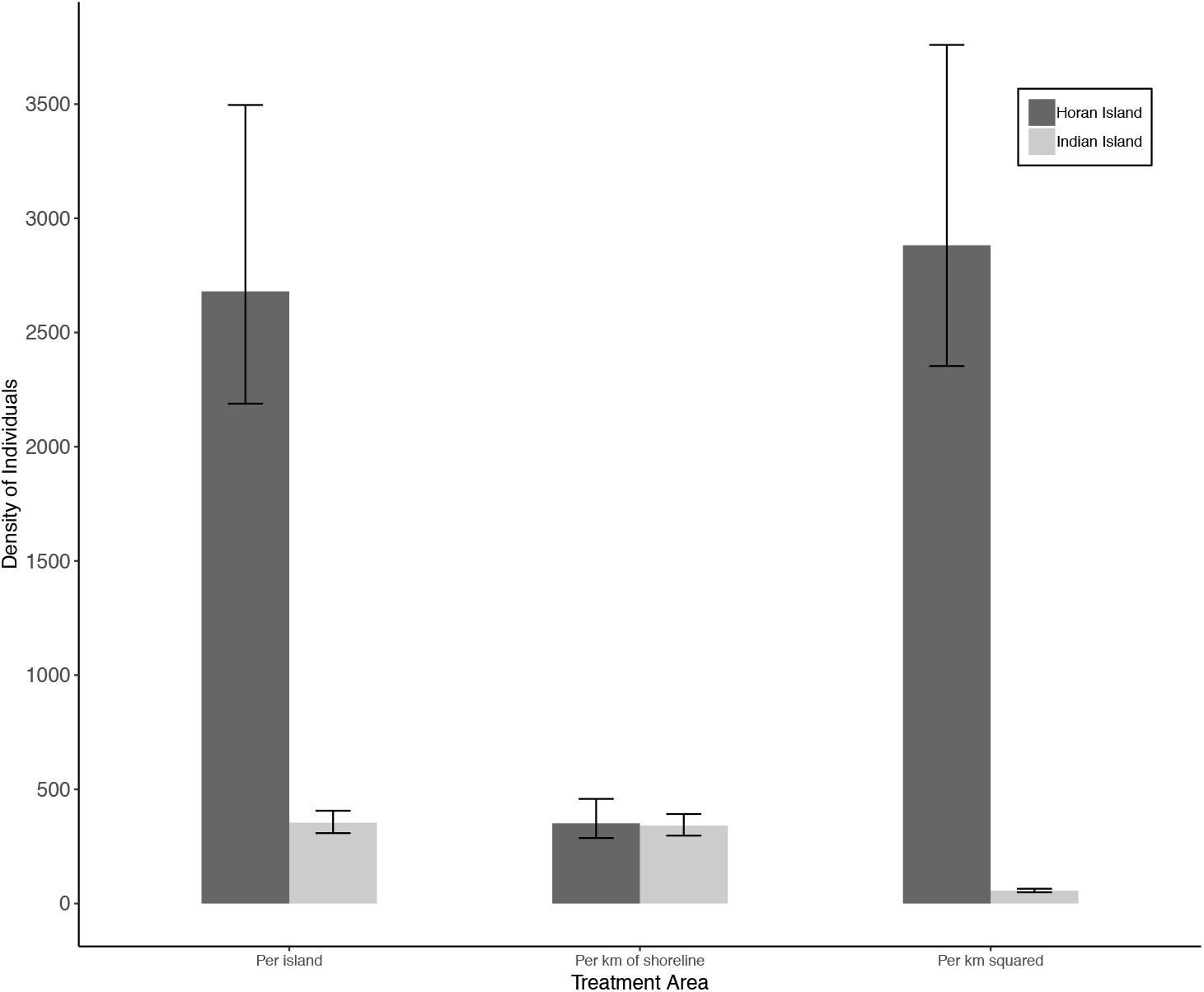
Estimated density of cane toads on each island using density calculated per island, per km, and per km^2^. There is clear concordance across islands when we calculate a linear density (per km).

Given the posterior estimates of *p* and *λ*, we examine the total survey effort (in days) required to eradicate toads on both Horan and Indian Island. Inputting the distribution of *N*_0_, leaving a single individual is equivalent to leaving the proportion, 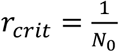 of the original individuals. The time to reach this point is given by ln(*r_crit_*)/ln(1 − *p*) = 75 days [49, 110] on HI, and 19 days [10, 29] on II (Figure 5).

**Figure 5.**
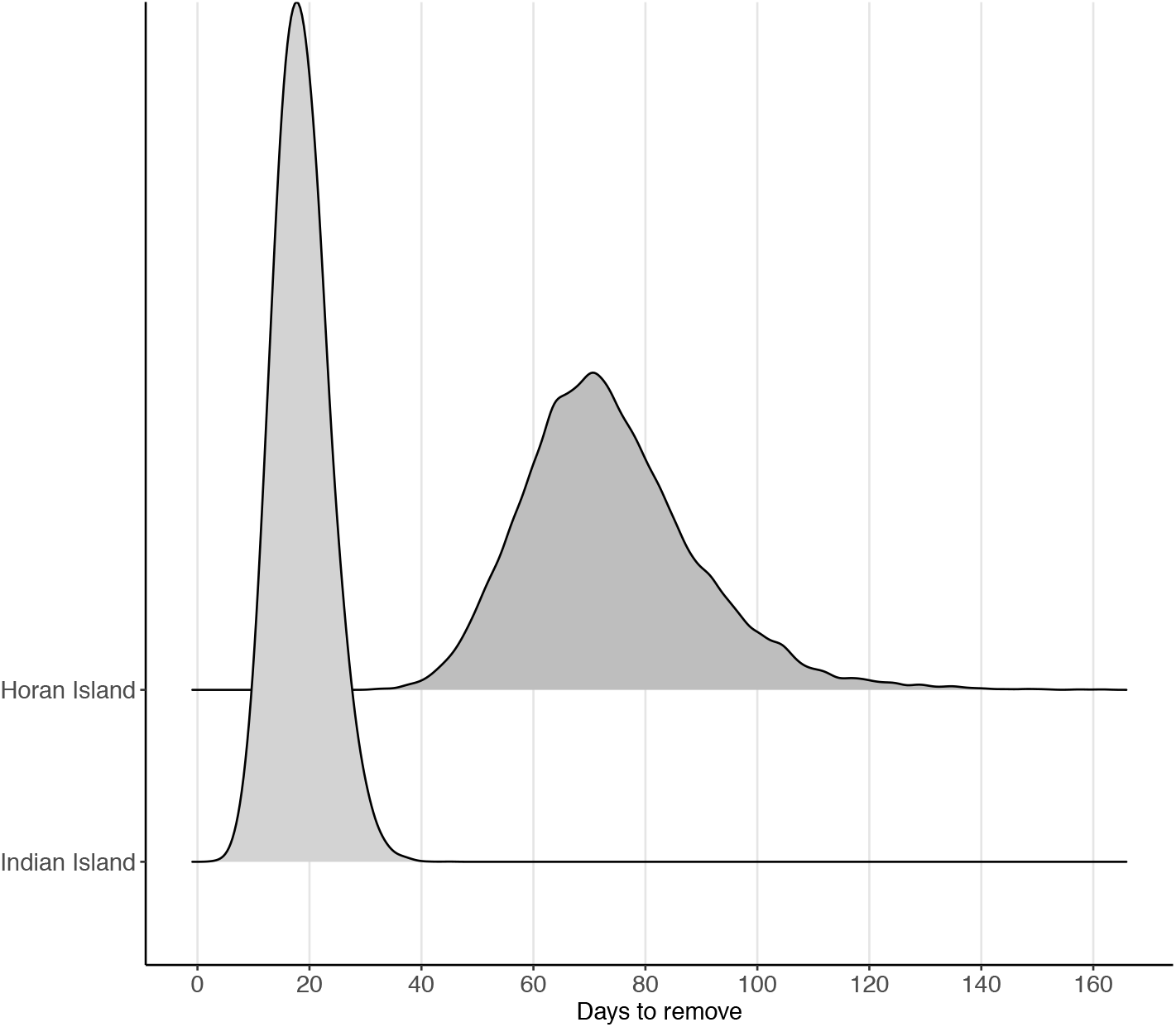
Distributions of the estimated numbers of days required to remove cane toads from Horan and Indian Islands.

### Cost Sensitivity

Multiplying the estimate of the number of days required to achieve eradication by our removal costs suggests that $62 407 [$40 583, $91 104] would be required to eradicate toads from HI. This equates to $9 748 [$6 339, $14 231] per kilometer of freshwater shoreline or $80 009 [$52 030, $116 801] per km$ of land. In contrast, the cost to eradicate toads from II is estimated to be $18 394 [$9 872, $28 636], equating to $17 737 [$9 523, $27 615] per kilometer of freshwater shoreline or $2 929 [$1 572, $4 560] per km$ of land (Figure 6).

**Figure 6.**
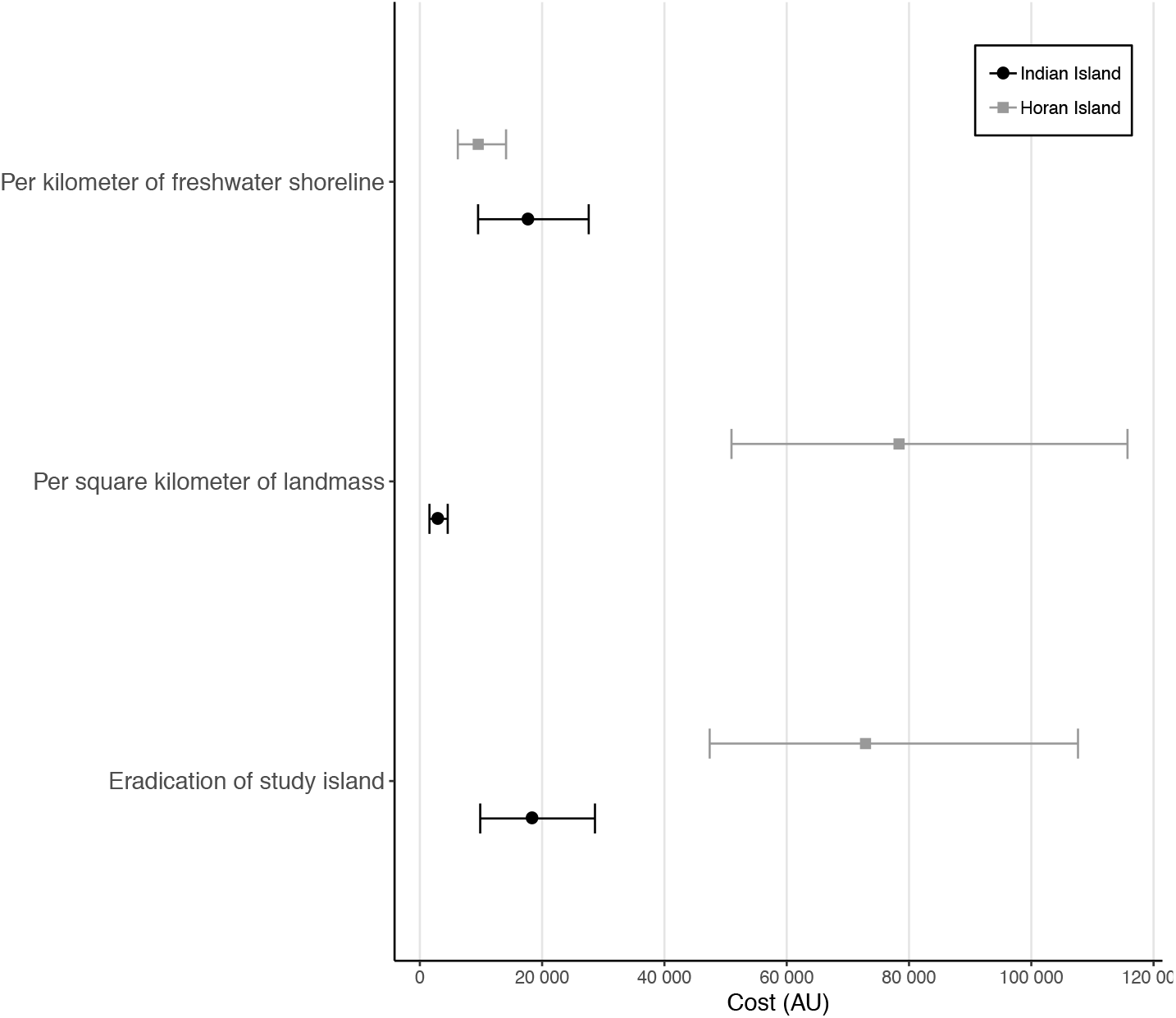
Costs of eradication calculated per km of shoreline, per square kilometre, and per island. Again, we see the strong concordance in costs when we calculate them per km of shoreline.

### Benefit of quarantine on Prioritized Australian Islands

Using our estimates of eradication costs per-kilometer of freshwater shoreline, we examine the economic benefit of cane toad quarantine on all toad-free islands (by jurisdiction), as well as the cost to restore all toad-inhabited islands to a toad-free state (Figure 7). The current economic benefit of quarantine on all prioritized toad-free islands is estimated to be between $17.5 M (HI estimates) and $33.3 M (based on II estimates). We estimate it would cost, on average, between $2.8 M (HI) and $5.2 M (II) to remove toads from all prioritized islands currently occupied by toads. Finally, we estimate the economic benefit of the ‘waterless barrier’ protecting the Pilbara to be between $34.3 M (HI) and $63.6 M (II).

**Figure 7.**
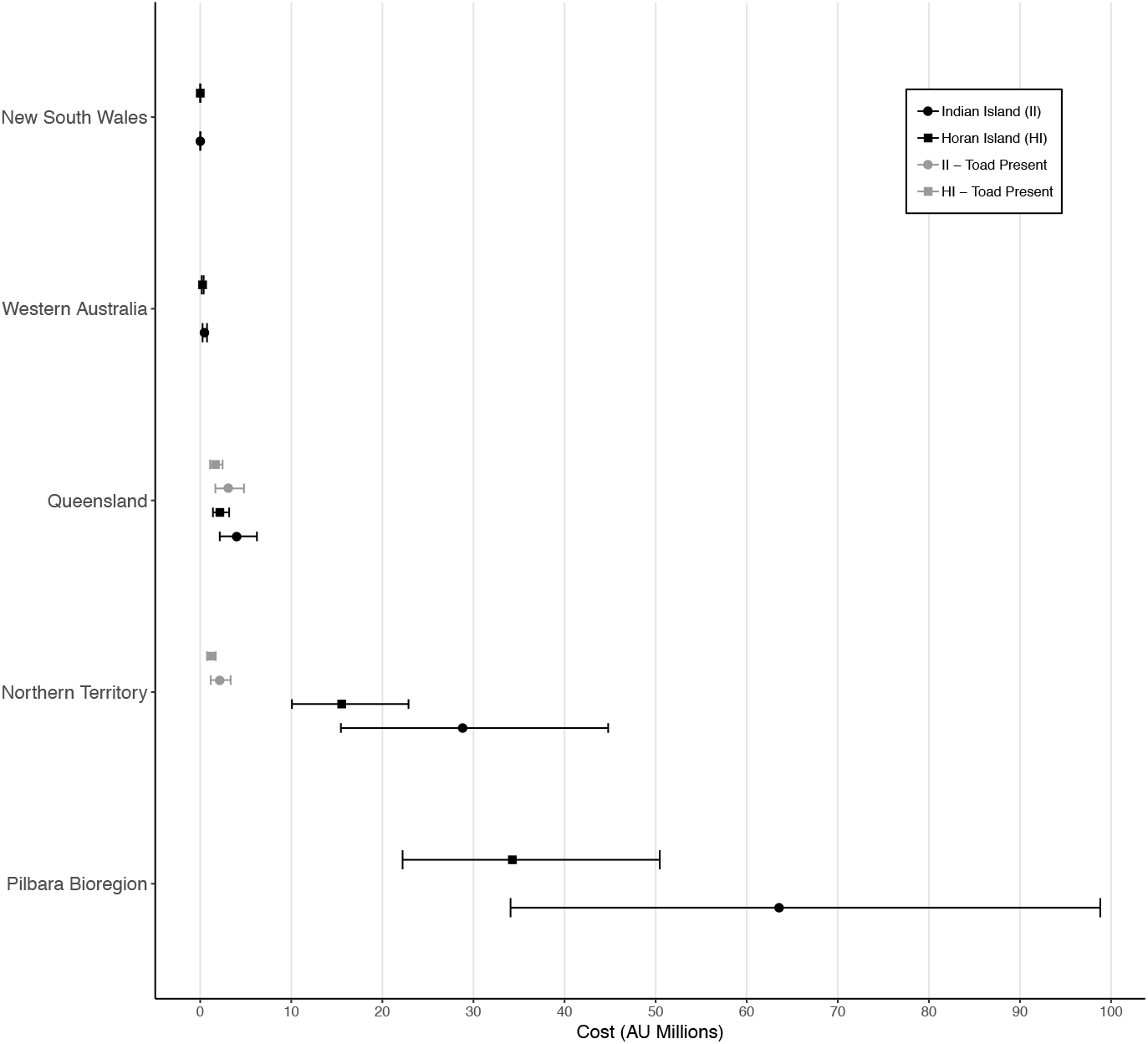
Distribution of the benefit of cane toad quarantine across different jurisdictions within Australia. Toad present distributions denote areas where toads are known to occur, and represent the cost to remove toads. No islands in either New South Wales, Western Australia or the Pilbara Bioregion have confirmed toad presence.

## Discussion

As the number of invasive species requiring management increases, practitioners must identify efficient strategies for allocating resources to various management activities. Although conventional wisdom places emphasis on prevention measures, the practice of valuing such actions in the face of non-economic costs can be challenging. Placing a monetary value on a conservation benefit will most often require some value judgement as to the monetary worth of biodiversity. Using estimates of a species detectability, population density, and subsequent eradication costs we aim to prevent such value judgement when investigating the benefit of quarantine measures in combatting the impact of the invasive cane toad across Australia’s prioritized offshore islands.

Despite substantial community and research effort into cane toad removal via trapping and hand capture, there are only a handful of published detection estimates for the species (Griffiths & McKay 2007). Our detection estimate is, of course, specific to the details of our survey. Nonetheless, it is surprisingly low for our large-shoreline site (Horan Island). Here, the length of shoreline meant we only passed each location once per night, and individual toads in this closed system had, on average, a 0.1 [0.07 – 0.13] chance of being seen on any given night. This contrasts with our small-shoreline site (Indian Island) where we were able to make multiple passes of the same point each night. Here, individual toads had a 0.27 [0.22 – 0.33] chance of being detected on a given survey night. Whilst individual toads are relatively easy to see when they are active, our results suggest that this might give a misleading impression of one-pass detectability.

We compared two density metrics: a linear density (per km) and an areal density (per km^2^). Our areal density estimate (2 852 individuals/ km^2^) is similar to estimates derived from previous studies of invasive cane toads in the Solomon Islands archipelago (1 035/km^2^; Pikacha *et al*. 2015), the islands of Papua New Guinea (3 000/km^2^; Zugg *et al*. 1975; Freeland *et al*. 1986), and density estimates of an analogous invasive toad on Madagascar (3 240/km^2^; Reardon *et al*. 2018). A single study conducted on the Australian mainland reported densities as high as 256 300 individuals per km^2^ (Cohen & Alford 1993), but this estimate was predominantly of the metamorph life stage, which occurs at very high densities prior to dispersal. Metamorphs are strongly constrained to the edges of water bodies (Child *et al*. 2008), and typically suffer high mortality from predation and desiccation before reaching maturity (Ward-Fear *et al*. 2010). While an areal density would make sense in a habitat where animals are constrained by some factor that scales with area (e.g., primary productivity), it is clear that toads in northern Australia are often constrained by access to water in the dry season, and thus length of shoreline is more appropriate. Length of shoreline not only defines access to water, but also the density of infectious parasites (such as *Rhabdias pseudosphaerocephala*) that use the moist conditions and high toad densities along shorelines as opportunities for transmission ((Kelehear *et al*. 2011; 2013). It is also likely that the survival rate of emergent metamorphs is dependent on length of shoreline, because this will set the density of conspecifics and so moderate the rate at which these conspecifics cannibalize each other (Pizzatto & Shine 2008). In comparing the areal and linear densities between our sites we find a large difference between sites in the areal metric, but a strikingly similar density value across sites in the linear metric. Our results suggest that across these two different systems, toads achieve a density of around 354 adults per kilometer of shoreline.

Because toads in dry conditions require regular re-hydration (Seebacher & Alford 2002; Tingley & Shine 2011) it is a logical step to conduct removal efforts when toads are restricted to a subset of semi-permanent hydration points during drier sections of the year (Letnic *et al*. 2015). Given the linear density metric is so concordant across sites, this is the best metric for calculating eradication costs. Certainly, if we use the areal metric we find a wide gulf in the possible eradication values in contrast to our shoreline metric (Figure 6). Costs based on the linear metric had a greater correspondence between the two sites ($9 748 and $ 17 737 AUD per km of shoreline). Encouragingly our cost estimates appear similar to estimates derived from a successful eradication program associated with removing the American bullfrog from two locations in Canada ($8 200 – $23 000 CAN per kilometer of freshwater shoreline).

To our knowledge, there is only one instance in which the cost to eradicate cane toads from an island has been documented (Wingate 2011). Carried out on Nonsuch Island in Bermuda, this removal occurred over six years and included countless volunteer hours and an investment of $10 000 USD (~$14 330 AUD) to remove toads from an area of 0.6 km^2^. In addition, two successful eradications from extralimital mainland sites have been documented, occurring beyond the southern border of the cane toads’ current range in Australia (White 2010; Greenlees *et al*. 2018). This handful of successful removals of the cane toad, mirrors a broad trend in the eradication of invasive amphibian populations globally (Adams & Pearl 2007; Kraus 2009; Beachy *et al*. 2011; Orchard 2011). As such, there is scant information available to guide policy makers and management agencies when evaluating the feasibility of implementing amphibian quarantine and eradication measures.

If we are to shift away from tactical, post-invasion approaches, to a preventative strategic approach, management practitioners require an estimate of the economic value that quarantine holds. Our analysis of the feasibility and benefit of cane toad quarantine is timely, given renewed emphasis on Australia’s offshore islands as safe-havens to buffer biodiversity against cane toad impacts. Sixty-two Australian offshore islands designated as ‘high conservation status’ fall within the cane toad’s predicted distribution; 21 of those have already been colonized by toads. Given our criteria (see Methods), we estimate the remaining value of toad quarantine across toad-free islands in northern Australia to be up to $33 M AUD. This value is conservative for a number of reasons. It is a reasonable expectation that as islands become home to increasing numbers of insurance populations or endangered species, the benefit of maintaining those islands as pest-free (measured as the cost of restoration) will increase. In addition, as toads establish onto an increasing number of these islands, those remaining toad-free will, by their scarcity alone, possess a greater economic and environmental value. While in reality it is unlikely that all islands without quarantine will be invaded, the benefit of quarantine within our dataset is held primarily in a few large islands (e.g. Melville Island, Table 1). These larger islands often have human settlements, competing management objectives (e.g., economic growth activities, multi-species quarantine) or more convoluted invasion pathways associated with anthropogenic activity. In short, quarantine needs to be carefully managed on these large islands. Eradication efforts for taxa other than toads have been successful on large islands (such as Santiago Island 5 465km^2^; Cruz *et al*. 2009), but they require much greater planning, intersectional management, and investment in post-eradication surveillance and monitoring (Moore *et al*. 2010, Rout *et al*. 2011, Carwardine *et al*. 2012). On these large islands, then, the value of prevention is very likely underestimated.

The vanguard of the cane toad invasion is currently sweeping across Western Australia at ~50 km per annum, but recent research suggests that a waterless barrier between the Kimberley and the Pilbara could halt the toad invasion (Florance et al. 2011; Tingley et al. 2013; Southwell et al. 2017). Applying our results to this management strategy revealed that the benefit of quarantine over such an area ($34.3 – $63.6 million) is roughly double the value of quarantine across all offshore islands combined ($20.3 – $38.5 million). The cost of quarantine in this case has been rigorously estimated at around $5 million dollars over 50 years (Southwell et al. 2017), only a fraction of what we estimate it would cost to eradicate toads from this area.

Here we demonstrate the immense benefit of toad quarantine across northern Australia. We avoid arbitrary judgement and simply calculate the cost of eradication in the case of quarantine failure. We acknowledge that this value is undoubtedly a lower boundary on the true benefit, but valuing preventative management is important. It becomes more so as conservation actions increasingly rely on offshore islands and fenced areas as cost-effective avenues to protect biodiversity from the impacts of invasive species. Quarantine measures often protect against multiple potential invaders but our results suggest that even when considering a single species, the monetary value of quarantine can be substantial. Prevention, it seems, is worth more than we might naively guess, even with aphorisms to remind us.

## Author’s Contributions

AS and BP contributed to all aspects of the manuscript. RT contributed to data collection and writing of the manuscript.

## Acknowledgments

We recognise and thank the Kenbi traditional Owners (Raylene and Zoe Singh) for land access permission. We thank Chris Jolly, Sarah McGoll-Nicolsen, John ‘ Mango’ Moreen and the Kenbi Ranger Group for their aid in the field, and for logistical support. Corrin Everitt, John Llewelyn, Ruchira Somaweera, and Greg Clarke provided constructive comments and advice. We also thank Greg Smith from Lake Argyle Cruises for his input and local knowledge. All procedures were approved by the University of Melbourne Animal Ethics Committee (1714277.1). This research was supported by an Australian Research Council Future Fellowship to BP (FT160100198) and an Australian Research Council DECRA to RT (DE170100601). Land access was granted via the Northern Land Council (permit 82368).

## Data Accessibility

Waterbody data are available upon request from Geographic Information Systems at the Department of Agriculture and Food, WA, and the Spatial Data Exchange section of the Department of water, WA. Cost data associated with our analyses can be found in the Supporting Information.

## Supporting Information

**Supporting Table 1:** Estimated costs of cane toad eradication derived from removal efforts in this study. All figures are in Australian Dollars ($AU).

| Study | Cost | Mea |  | Low | Uppe |
| --- | --- | --- | --- | --- | --- |
| Horan | Consumable | 6 330 | ± | 4 | 9 320 |
|  | Personnel | 50 | ± | 32 | 74 |
|  | Travel | 16 | ± | 10 | 24 |
| Indian | Consumable | 1 595 | ± | 855 | 2 480 |
|  | Personnel | 12 | ± | 6 | 19 |
|  | Travel | 4 142 | ± | 2 | 6 438 |

